# Evidence for adaptive evolution towards high magnetic sensitivity of potential magnetoreceptor in songbirds

**DOI:** 10.1101/2023.03.25.534197

**Authors:** Corinna Langebrake, Georg Manthey, Anders Frederiksen, Juan S. Lugo Ramos, Julien Y. Dutheil, Raisa Chetverikova, Ilia Solov’yov, Henrik Mouritsen, Miriam Liedvogel

## Abstract

Migratory birds possess remarkable accuracy in orientation and navigation, which involves various compass systems including the magnetic compass. Identifying the primary magnetosensor remains a fundamental open question. Cryptochromes (Cry) have been shown to be magnetically sensitive, specifically Cry4 shows enhanced magnetic sensitivity in migratory songbirds compared to resident species. Here, we investigate cryptochromes and their potential involvement in magnetoreception in a phylogenetic framework, integrating molecular evolutionary analyses with protein dynamics modeling. We base our analysis on 363 bird genomes and associate different selection regimes with migratory behaviour. We show that Cry4 is characterized by strong positive selection and high variability, typical characteristics of sensor proteins. We identify key sites that likely facilitated the evolution of a highly optimized sensory protein for night time compass orientation in songbirds and a potential functional shift or specialisation. Additionally, we show that Cry4 was lost in hummingbirds, parrots and Tyranni (Suboscines) and thus identified a natural comparative gene knockout, which can be used to test the function of Cry4 in birds. In contrast, the other two cryptochromes Cry1 and Cry2, were highly conserved in all species, indicating basal, non-sensory functions. Our results strengthen the hypothesised role of Cry4 as sensor protein in (night)-migratory songbirds.

## Introduction

Juvenile night-migratory songbirds on their first migration fly at night on their own to an zarea they have never been before, which emphasizes that their ability to orient accurately over long distances is considered a key adaptation for migration [1]. Migration requires remarkable orientation and navigation abilities combined with inherited information on timing and direction, which are integrated into a spatiotemporal orientation program that allows birds to migrate between breeding and non-breeding grounds with extraordinary precision [2-4]. It is therefore not surprising that seasonal bird migration is characterized by a number of seemingly associated traits, including the propensity to migrate, circannual timing, physiological adaptations such as fattening, as well as the ability to determine the distance and direction of migratory movement [1,5].

Evolutionary scenarios assume that these migratory traits are heritable. Artificial selection and crossbreeding studies showed that several migratory traits have a major genetic component, and can drastically change within a few generations when under strong artificial selection [6-9]. Especially the inheritance of migratory direction observed in crossbred European blackcaps *Sylvia atricapilla* suggests that the inheritance of migratory direction is governed by variations in a small number of genes [7-8].

Evolutionary scenarios also assume that migratory behavior is evolving on a phylogenetically relevant timescale. At first glance, it is not obvious that this assumption is actually met. The migratory phenotype can be very diverse between closely related species and sometimes even varies within different populations of the same bird species [9-11]. However, while it is clear that the way migratory birds use their fundamental sensory capabilities to generate a migration strategy (e.g. how different cues are calibrated, what direction birds migrate in, and whether a specific population or individuum is migratory or not) can change very quickly, the essential underlying adaptations for migration, such as the sensory capabilities (e.g. the ability to sense a magnetic field or the ability to use the moving pattern of the stars) are assumed to evolve on a much slower time scale [1,12,13]. For example, most night-migratory songbirds drastically change their normally strictly diurnal behavior to a migratory state which includes long-distance nocturnal flights, but day-migrants do not seem to be able to change into night-migrants [1]. Piersma et al. (2005) [1] therefore conclude that night-time compass orientation and the sensory capabilities required for it are migratory adaptations, which might be deeply anchored in the avian phylogeny.

To determine their migratory direction, night-migratory birds possess three different compasses: a sun compass, a star compass, and a magnetic compass [3,14,15]. The geomagnetic field is an omnipresent reference system and of major importance for successful navigation [3]. Therefore, the genes coding for the molecules involved in the magnetosensory pathway could be under particularly strong selection pressure and would be especially exciting to study from an evolutionary perspective on a phylogenetic scale.

Various mechanisms for magnetoreception have been proposed, but increasing evidence suggests that a light-induced radical pair-based mechanism in cryptochrome (Cry) proteins form the basis for magnetoreception in night-migratory songbirds [16-19]. Cryptochromes are blue light photoreceptors usually discussed in the context of circadian rhythm regulation. They are the only known photoreceptor molecules in the bird’s eye that, at least in some cases, can use light energy to form long-lived, magnetically sensitive, radical pairs [18-20]. Three different cryptochrome genes have been identified in birds, namely Cry1, Cry2 and Cry4, each with two splicing variants, referred to as a and b variants respectively (reviewed in [21]) [22-23]. Cryptochrome 4a (Cry4a), which has no other described functions next to magnetoreception, has received the most attention as a possible magnetosensory protein in birds as, in contrast to Cry1 and Cry2, Cry4 is known to bind flavin adenine dinucleotide (FAD), the crucial chromophore co-factor for radical pair formation in cryptochromes [19,24-27]. Cry4a is expressed in double cones [26] that form mosaics within the retina of several bird species which could aid magnetoreception [28,29]. Furthermore, while Cry1, Cry2 and Cry4b show clear circadian changes in expression, Cry4a is stably expressed over a 24h period in birds, but shows increased circannual expression levels during the migratory seasons [22,26,30]. In fish, Cry4(a) does show a circadian oscillation pattern over a 24h period, but this is absent in all bird species investigated so far [22,26,30,31], suggesting a functional change between the two clades. Magnetically sensitive radical pairs form between FAD and a chain of four tryptophans (Trp) in Cry4a [19]. Importantly, it has been suggested that these radicals in Cry4a of night-migratory songbirds show increased magnetic sensitivity compared to the equivalent Cry4a radical-pairs from non-migratory species, suggesting an optimization of magnetosensing in night-migratory songbirds [19].

Based on the recent publication of a large number of high-quality avian reference genomes [32], it is now possible to approach this fascinating topic of magnetoreception from a novel angle using a phylogenetic approach. Therefore, the aim of the present study is to characterize similarities, variation and deletion of coding DNA sequences between different members of the cryptochrome multigene family across the avian clade. We hypothesize that a novel and strikingly different function of a cryptochrome-derived magnetosensor will be reflected by a distinct evolutionary history compared to cryptochrome members with an essential role in circadian rhythm regulation that is highly conserved across most vertebrates [33-35]. Specifically, we predict that a protein with a highly conserved function in different taxa would show very little variation in its amino acid composition across the avian clade. In contrast, a protein that evolved an additional sensory function such as magnetoreception within the avian clade, should be characterized by increased variability and positive selection on several key amino acid residues essential for this novel/optimized sensory function of the specific cryptochrome. We explore different scenarios of selection and functional changes across taxonomical and behavioral groups to pinpoint sites where evolution may have optimized and shaped a putatively magnetosensory protein.

## Results

### Cry1 and Cry2 are highly conserved and present in all birds, while Cry4 shows high variability and complex losses in three clades

Cry1 and Cry2 nucleotide sequences were extracted for all species and were characterized by high sequence conservation. Only few bird species showed higher fragmentation with exons missing, though the first exon was incomplete in some species (for details see Supplementary Data S1).

In strong contrast, Cry4 sequences varied much more between species, in both the photolyase and FAD-binding domains (Figure 1). Because of this clear difference and the suggested role of Cry4 as a magnetosensory molecule, we investigated whether Cry4 is actually present in all birds or might have been lost in certain, possibly resident, clades. This is a pattern observed in other sensory proteins, such as the opsin family, where complicated patterns of duplications and losses associated with species ecology have been described [36-38]. Our analyses suggest the apparent loss of Cry4 in at least three clades with mostly resident tropical species: parrots (Psittaciformes), hummingbirds (Trochilidae) and a group of Passeriformes, the Tyranni (Suboscines) (Figure 2). Except for apparent losses of the gene in these three clades, we could extract complete Cry4 sequences for most other birds including migratory, resident and even flightless birds (for details see Supplementary Data S1).

**Figure 1:**
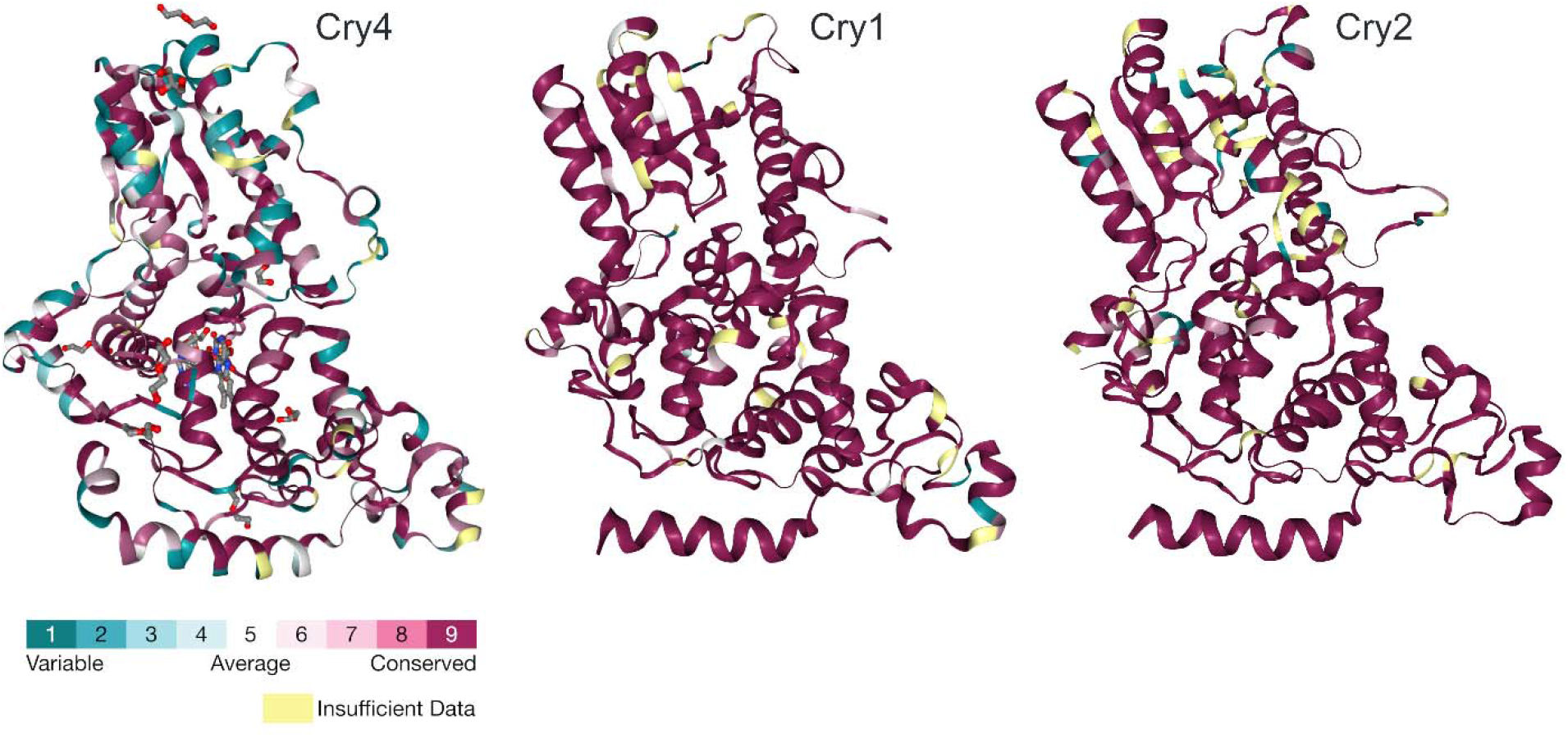
Variability in the amino acid sequence of different cryptochromes across Aves. Cryptochrome 4 (Cry4, Reference Sequence 6PTZ from the RSCB Protein Data Bank) is characterized by a high amount of variability between different bird species, in both the photolyase and FAD-binding domains, whereas Cry1 (6KX4) and Cry2 (7D0N) are strongly conserved. The C-terminus which shows high variability in all three proteins and which has not been crystalized is not displayed. Purple = conserved, green = variable, yellow = unknown due to insufficient data. The conservation scores (dimensionless) are normalized so that the average score for all sites within a protein equal zero. Unknown (yellow) classification results from sites in the alignment where less than six species provide reliable aa information or where the confidence interval around the calculated score is spanning 4 or more conservation classes (graphs generated by ConSurf; Ashkenazy et al., 2016, 2010; Landau et al., 2005).

**Figure 2:**
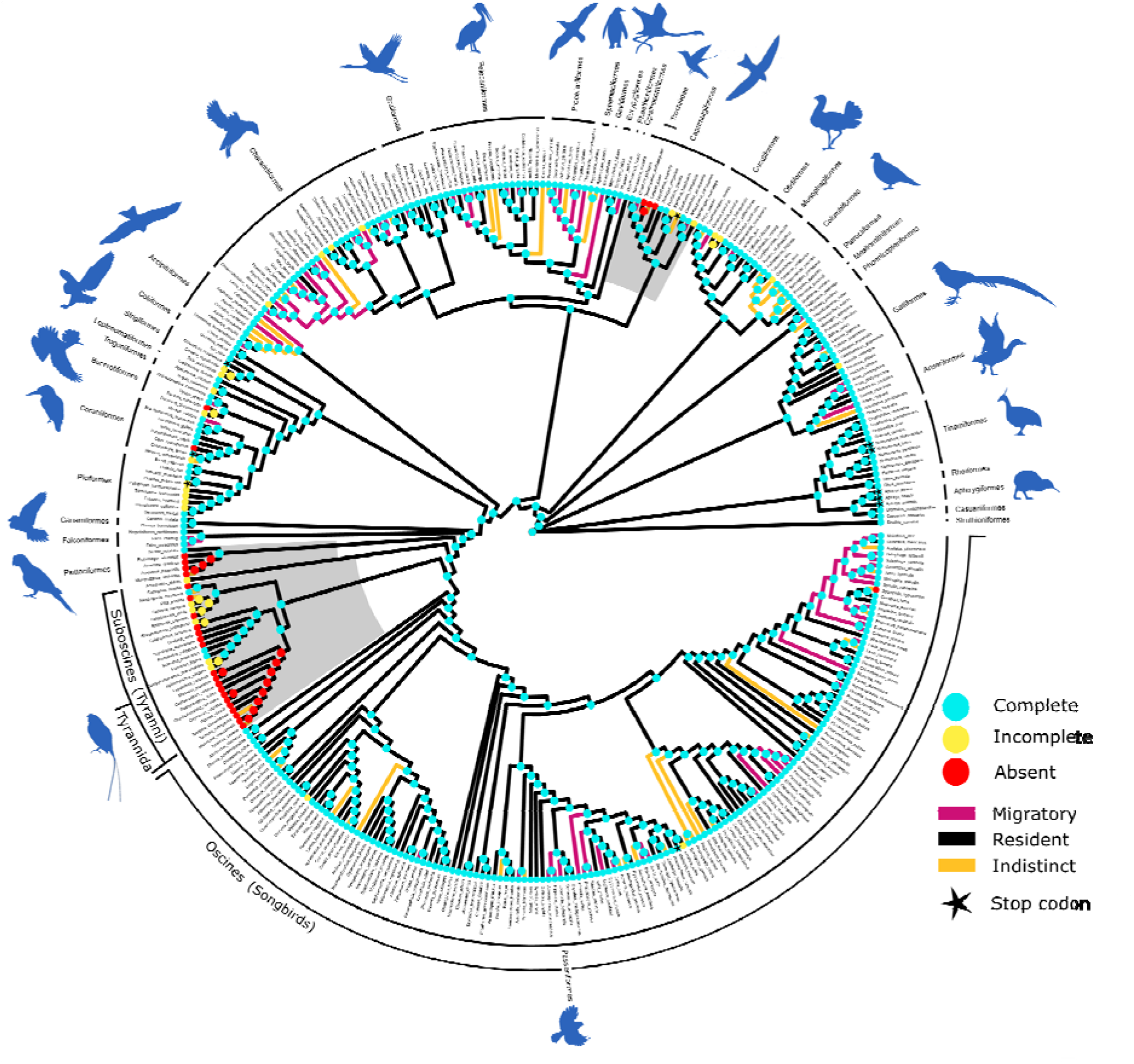
Three independent Cry4 losses occurred across the avian phylogeny. Cry4 was completely lost in three clades (red nodes): hummingbirds, parrots and Tyranni (Suboscines), highlighted in grey inlet, whereas most other species retain a complete Cry4 sequence (turquois nodes). Species with incomplete Cry4 sequences are indicated by yellow nodes. Cry4 was mapped onto the B10K phylogeny. Species are characterized with respect to migratory behavior as indicated by the branch color: black = resident, purple = migratory, and orange = indistinct (no clear phenotype characterization possible) behavior. If any stop codons occur in the coding sequence, this is indicated by a star. Seemingly isolated absences of Cry4 (red dot at the species level) in single species are likely due to fragmented assemblies.

In the context of magnetoreception and magnetic compass orientation, the apparent loss of Cry4 is particularly interesting in the Tyranni. The long sedentary and tropical history of most families in the Tyrannida including an apparent failure to evolve a migratory lifestyle in some families [13,39] and the loss of a putatively magnetosensory protein could be related. However, this family now includes several long-distance, night-migratory species, such as the *Empidonax* genus that evolved recently (late Pliocene/early Pleistocene, 3-2 million years ago (mya); [40,41]). To further investigate the loss of Cry4 in Tyranni, we characterized several different states of fragmentation of the Cry4 protein within the Tyranni, but could assign a true loss to the first occurrence of Tyrannida (Figure 2). This loss was confirmed by synteny analyses between the high-quality genome of the lance-tailed manakin *Chiroxiphia lanceolata* (Pipridae within the Tyrannida, GCA_009829145.1) and the Eurasian blackcap *Sylvia atricapilla* (GCA_009819655.1), another high-quality reference genome within songbirds. The Cry4 gene region covering ∼20.000 base pairs (bp) in the Eurasian blackcap was reduced to roughly 5.000 bp in the manakin without any traces of Cry4 genomic sequence left (Figure 3). Our results suggest that Cry4 was first subject to pseudogenisation as indicated by the ancestral reconstruction showing old world Tyranni species (such as the common sunbird-asity *Neodrepanis coruscans*) that still have gene fragments left, followed by gene loss in the Tyrannida.

**Figure 3:**
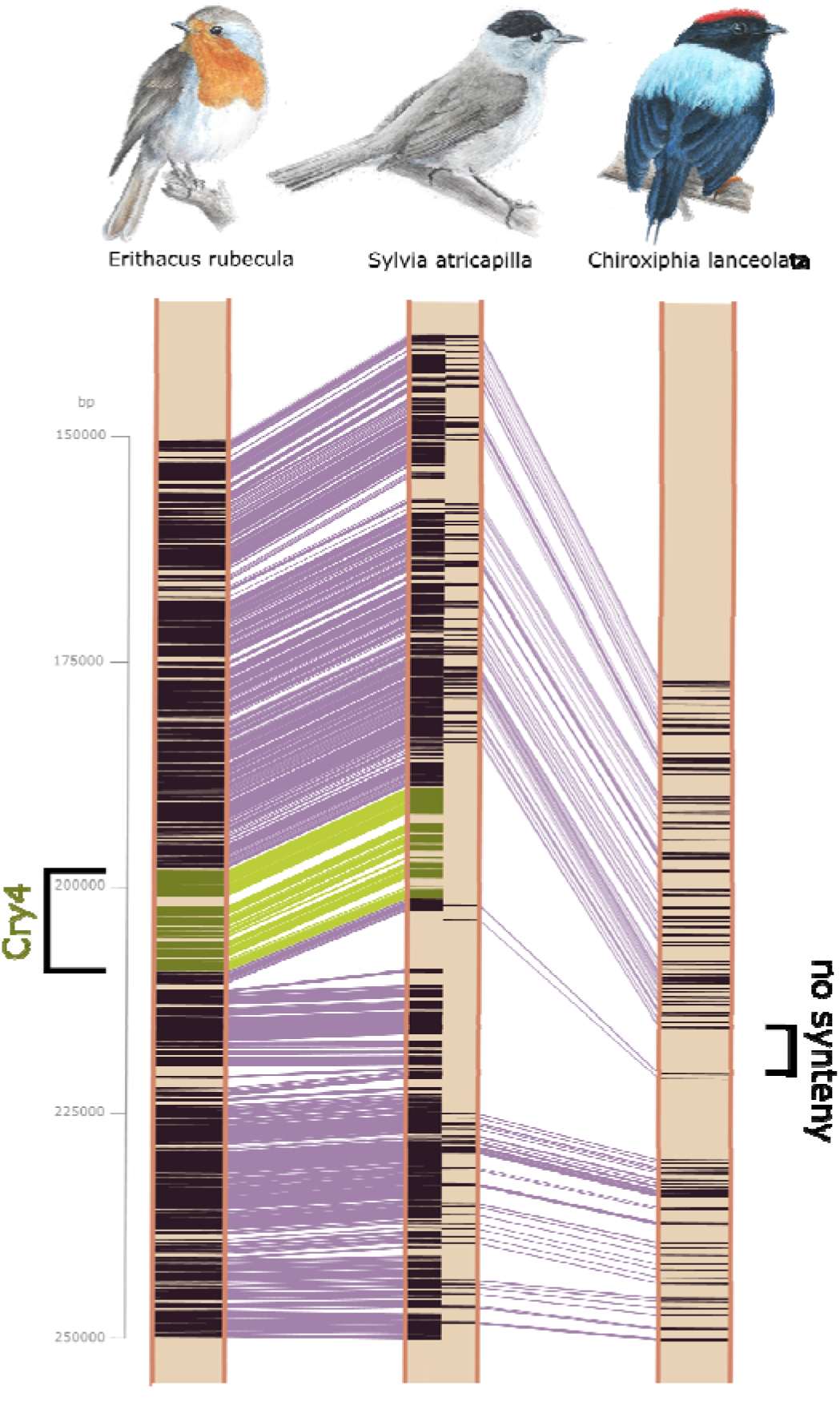
Synteny analyses reveal deletion of Cry4 in Tyranni. Synteny analyses between the high-quality genome of the lance-tailed manakin (Chiroxiphia lanceolata; Pipridae within the Tyrannida) and two songbird species, the European robin (Erithacus rubecula) and the Eurasian blackcap (Sylvia atricapilla) clearly show that the Cry4 gene region covering ∼20.000 base pairs (bp) in both songbird species (green) was reduced to roughly 5000 bp in the lance tailed manakin (Chiroxiphia lanceolate) (right) without any traces of the Cry4 genomic sequence present. Green = Cry4 synteny, purple = other regions of synteny.

To prove the absence of Cry4 in long-distance night migratory representatives within the tyrant flycatcher, we used RNAseq to assemble a high-quality reference-free transcriptome based on tissue from brain, retina and ovaries for the least flycatcher (*Empidonax minimus*) (BUSCO: 91.2 % complete genes; realignment rate with bowtie 96.9 %), and we added the reference genome of the willow flycatcher (*Empidonax traillii*, PRJNA437496) to complement our B10K dataset [42]. In contrast to Cry1 and Cry2, Cry4 was absent from both Empidonax flycatcher’s genome and transcriptome. These results conclusively confirm the absence of Cry4 in *Empidonax* flycatchers.

In parrots, Cry4 is still present in the New Zealand species (Kea and Kakapo, GCA_004027225.2), but was lost in all other species (Figure 2). To investigate the loss of Cry4 in hummingbirds, we used an available dataset for a liver transcriptome of the ruby-throated hummingbird *Archilochus colubris* [43] (PRJNA369257) to characterize the presence or absence of all cryptochrome members and detected transcripts for both Cry1 and Cry2 whereas Cry4 was absent, further corroborating the loss of Cry4 in hummingbirds.

Although the loss of Cry4 does not seem to be linked to residency, the loss of the extra exon in Cry4b (11 instead of 10 exons in Cry4a) through deletion or nonsense mutations in songbirds is significantly associated with migratory phenotype, suggesting a functional disadvantage of the extra exon in Cry4b compared to Cry4a in migratory songbirds (Figure S1, S2, LRT: 15.37, df: 3, p-value: 0.0015). The D-statistic [44] revealed a strong phylogenetic signal for both traits (migratory/resident phenotype: D = -0.082; extra exon present/absent: D = 0.118) justifying the correction for phylogenetic background when testing the association of both traits.

### Cry4 is characterized by positive selection and shows shifts of selection pressure in passerines

Xu et al. (2021) [19] identified a higher magnetic sensitivity of Cry4 in a night migratory songbird compared to resident chicken and pigeon, possibly reflecting adaptive evolution. Thus, we test how and where evolution optimized and shaped Cry4 as a putative magnetosensory molecule within the avian clade.

To detect patterns of positive diversifying selection, we used the codeml site-model and branch-site model [45,46]. Specifically, for the branch-site model we sequentially selected (i) passerines, (ii) migratory passerine clades, as well as (iii) branches leading to the passerines, (iv) migratory clades and (v) the non-passerines as foreground branches.

The likelihood ratio test (LRT) for the nested site-models of Cry1a and Cry1b was not significant and characterized Cry1a and b by strong negative selection; no sites in Cry1a or b were under positive selection. Similarly, Cry2 was characterized by negative selection, except for two amino acids sites (residue 569 and 583) located at the C-terminus which were identified by the Bayes Empirical Bayes (BEB) analysis to be under positive selection (Table S3).

For Cry4a, seven amino acid sites were identified with both site-model comparisons (the LRT was significant for M1a - M2a and M7 - M8 comparison) to be under positive selection with the BEB analysis (Figure 4a, Table S1/S2). The following four positively selected sites (residue 83, 68, 116, 189) are surface exposed and placed in a strikingly symmetrical non-random pattern around the protein. We also investigated the planar nonrandom position of the four residues by modeling a plane through all four sites and simulating its stability over time, confirming its nonrandom configuration (Figure 4a, see supplementary methods and results for more information).

**Figure 4:**
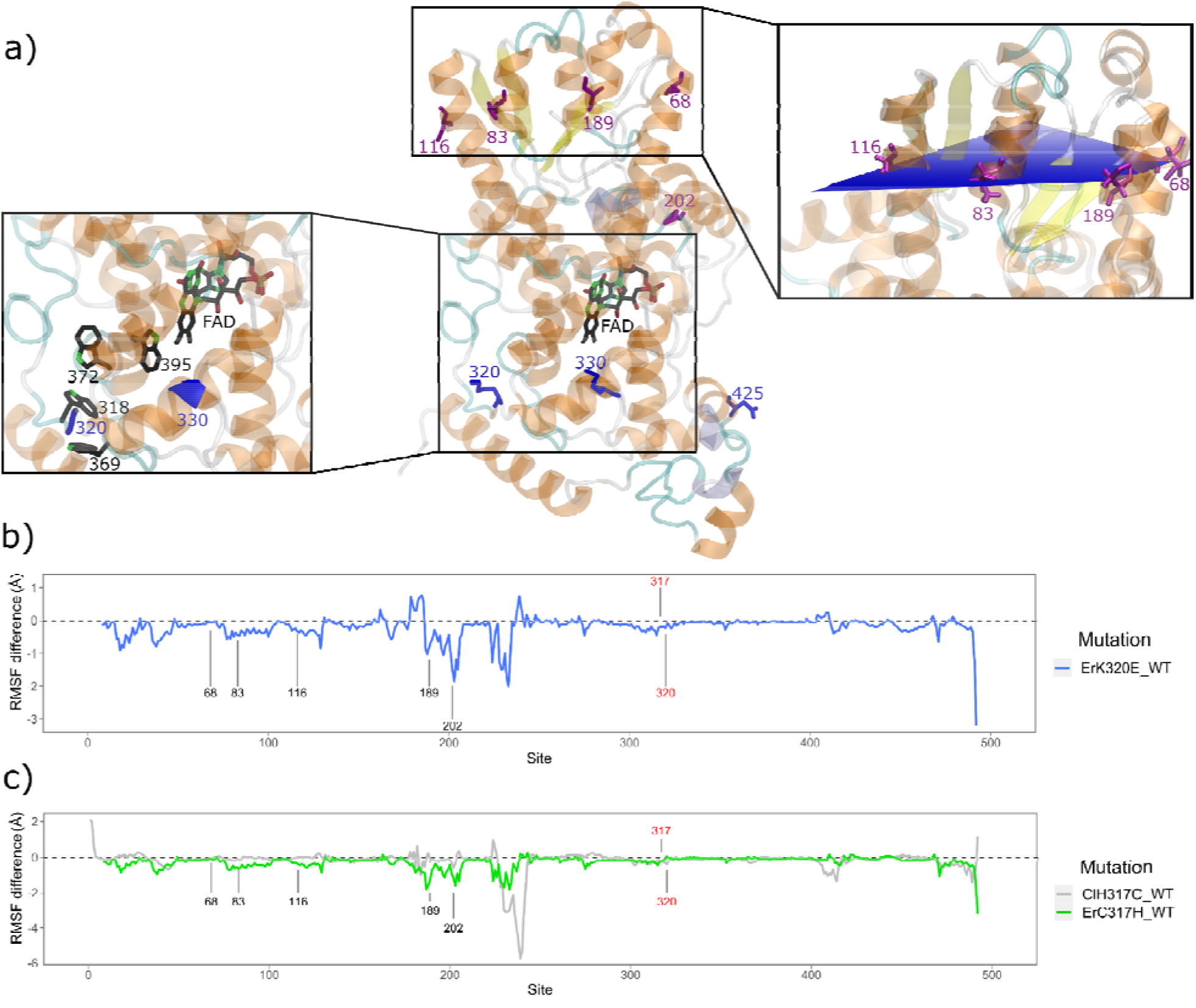
Sites under positive selection in Cry4 and site fluctuation changes after targeted mutations at diagnostic sites. **a)** ErCry4 (middle) showing the FAD chromophore and sites under positive selection identified in all species (purple) and only in passerines (blue). Notice that all positive selection near the FAD – tryptophan tetrad only occur in passerines. Magnified inlets to the left and right highlight detailed locations of the positively selected sites in passerines relative to the tryptophan-tetrad (318, 369, 372, 395; left) and the symmetrical position of four residues under positive selection across all birds (68, 83, 116, 189; right). **b)** The amino acid (aa) lysine (K) found at the positively selected site 320 in the European robin Cry4 was computationally mutated into the aa glutamate (E) found at site 320 in resident pigeons Cry4 (ErK320E), and the blue line shows the resulting change in protein fluctuation calculated as Root Mean Square Fluctuation (RMSF, in Ångström). Peak fluctuation changes occurred close to candidate sites 202 and 189 (see a). The protein always has two highly flexible domains: the phosphate binding loop and C-Terminus, which are mostly not resolved in the crystal structure (residues 228-244, 497-527). **c)** Site 317 showed an evolutionary rate shift towards higher conservation in migratory passerines. We therefore computed changes in protein fluctuations for ErCry4 with site 317 (cysteine, C, in robin) mutated into the resident pigeon correspondent (histidine, H). Fluctuations for ErC317H are shown in green and highlight peak fluctuation changes specifically at candidate sites 202 and 189. As a complementary test, we mutated the aa at site 317 of the pigeon Cry4 (ClCry4) into the robin aa (ClH317C), the resulting protein fluctuations are shown in gray.

We excluded Cry1 and Cry2 from further investigations with the branch-site model, because PAML struggled to converge due to high sequence conservation.

For Cry4a, the likelihood ratio test of the branch-site model was only significant when passerines were compared to all other species. Following the BEB analysis, seven sites were identified to be under positive selection in passerines (Figure 4a, Table S1/S2). Two of these seven sites (residue 202 and 510), were also identified as under positive selection in the whole phylogeny. The evolutionary rate at amino acid position 202 was constant in all species, but at position 510, it was significantly higher in passerines compared to non-passerines (tested with rateshift, see below) and a Contrast-FEL analysis identified divergent selection pressure at site 510 (table S5). Therefore, we conclude that the strong selection signal from the passerines was driving the significant result in the site-model, and suggest that amino acid residue 510 evolved neutrally in non-passerines, which the branch-site model also confirmed. Amino acid sites 330, 425, 512 and 525 were characterized by neutral evolution in the site-model, suggesting a shift from neutrality to positive selection in the passerines, which is supported by the results of the branch-site model. Site 320 was under positive selection only in the passerines, but was conserved and thus under negative selection in non-passerines. This distinct pattern indicates a major functional shift of this particular amino acid site 320 in passerines, mutating from the negatively charged glutamic acid to the positively charged lysine. Both amino acid site 320 and 330 are located within the potentially magnetosensitive area and are positioned close to the tryptophan tetrad of Cry4 (Figure 4a).

The comparison of non-passerines as the foreground clade to passerines was not significant, indicating that no sites were under positive selection in non-passerines.

The analysis with Hyphy [47] FEL and Contras-FEL supported our analyses with PAML (see supplementary results and Table S4 and S5). Especially residues 320 and 510 are detected recurrently. Another approach to detect directional selection was also able to retrieve candidate sites identified with PAML (residues 116, 202, 516 and 525, see supplementary methods).

### Evolutionary rate shifts associated with the migratory phenotype in passerines

Shifts in evolutionary rate of a certain site can indicate functional shifts of that focal site and allow us to link migratory behavior to rate changes at each site in the protein. Specifically, we were interested in sites that showed a higher conservation in migratory passerines compared to other species. As the substitution rate depends on the selection regime, at constant mutation rate, a substitution rate shift represents a signature of change in selective constraints. A neutral site acquiring a new functional role will see its substitution rate decreasing, while a relaxation of the selective pressure at a functional site will lead to an increase in its substitution rate. Alternatively, recurrent episodes of adaptive evolution may also lead to an apparent increased substitution rate. The comparison of Cry4 in passerines and non-passerines revealed 51 amino acid sites with a significant rate shift (ca 10 % of all sites, see Supplementary Data S2). Two sites experienced a particularly strong shift: amino acid residue 31 shifted from less constrained evolution in non-passerines to high conservation and no variability on the amino acid level in passerines, whereas residue 510 shifted from a low evolutionary rate in non-passerines to a high evolutionary rate in passerines, possibly reflecting the shift to positive selection detected by the branch-site model. The comparison of passerine clades with a migratory ancestor to the rest of the tree identified 18 sites that experienced a rate shift. Four amino acid sites (residues 45, 89, 201, 317), show a lower evolutionary rate in migratory clades along with higher conservation (Table S1 and Figure S4). Amino acid residue 89 experienced an especially strong shift from the previously dominant valine to isoleucine, which is not variable in migratory passerines. This site is surface exposed and located on top of the four symmetrically ordered amino acids that we characterized under positive selection (Figure S4). Amino acid site 317 should be highlighted as a site where all migrants show extreme conservation and exclusively exhibit a cysteine. Furthermore, the cysteine at residue 317 is only present in passerines. Residue 317 is located directly in the magnetosensitive area of the protein and a direct neighbor to the tryptophan at site 318 (TrpC), involved in the electron transfer chain of the radical pair forming process [19]. None of the identified sites with a special focus on the four symmetrical sites showed a signature of co-evolution (Figure S3 and supplementary methods).

To further quantify the impact of changes in the local structure of the protein near the electron transport chain of Cry4 on the overall structure of Cry4, we have performed molecular dynamics simulations of the European robin Cry4 (*Erithacus rubecula*, ErCry4) [19,26,48] and mutated the wild type protein at key candidate sites in the conserved tryptophan tetrad (site 317 and 320) into the pigeon (*Columba livia*) state, which is a resident species. Specifically, the cysteine (C) residue 317 in the ErCry4 structure was mutated into histidine (H, ErC317H), and the lysine (K) residue 320 was mutated to glutamate (E, ErK320E). Furthermore, residue 317 in the pigeon Cry4 (ClCry4) was mutated conversely into cysteine to mimic the robin state (ClH317C). Results show, that the mutations at both sites in ErCry4 impact the fluctuation of restricted regions in the whole protein. Especially strong shifts towards a lower fluctuation and thus a higher stability occur around the sites 202 and 189 (Figure 4b and c), which were also identified under positive selection. The independent molecular dynamics analysis thus suggests that the candidate residues in the magnetosensitive area might interact with sites which appear under positive selection in the photolyase domain far from the tryptophan-tetrad. The fluctuation changes in the ClCry4 mutant were overall small and not as pronounced as those in the ErCry4 mutants (see supplementary results for more information).

## Discussion

High evolvability driven by sites under positive diversifying or directional selection has been shown to aid niche adaptation in sensory proteins [36,38,49-54]. By combining molecular evolutionary analysis and protein dynamics simulations, we identify candidate sites that could help us to understand Cry4 functionality. Cry4 is characterized by high sequence variability and positive selection, and certain amino acids at specific sites correlated with migratory behavior. In contrast, Cry1 and Cry2 show extreme sequence conservation and almost no variability. This pronounced difference is in line with previous analyses of cryptochrome evolution [31,33,34,35,55,56] and functionality [19,26,29,55,57] and corroborates the suggestion that Cry4 fulfills at least one distinct sensory function, whereas Cry1 and Cry2 primarily seem to be circadian clock proteins. Furthermore, evolutionary scenarios suggest that genetic features associated with migratory traits evolved from common genes that then specialized further in migrants [1]. The recent findings by [19] suggesting that Cry4 from night migratory species might have increased magnetic sensitivity compared to non-migratory (resident) chicken and pigeon strengthen this hypothesis.

Based on our current knowledge [3,15,18], a cryptochrome-based magnetic compass receptor has to comply with several key requirements: (i) It needs to be magnetically sensitive, which requires binding of the co-factor FAD, (ii) it must stably bind to a surface to work as a compass, and (iii) it must initiate a biochemical amplification cascade [18]. In our phylogenetic analyses at the sequence level, we find several changes that could be involved in fulfilling these requirements.

In Cry4, almost 10% of all amino acid sites in the protein significantly changed their evolutionary rate from non-passerines towards passerines, and, specifically in passerines, key sites experienced a shift to positive selection. We furthermore observed that some of the amino acids that changed in passerines got subsequently “fixed” (show significantly reduced evolutionary rate or higher fitness) in migratory passerines. Such drops in evolutionary rate following a substitution typically indicate conservation of successful changes in a protein’s function through directional selection. Thus, we suggest that Cry4 might have experienced a main functional shift or specialization in passerines, potentially connected to magnetosensing, which is in line with recent spectroscopic findings studying Cry4 *in vitro* [19]. In particular, the residues 320 and 317 are located near the conserved tryptophan chain of the FAD-binding domain and could very likely directly influence radical pair formation or stabilization and potentially mediate the suggested elevated magnetic sensitivity of Cry4 in night migratory songbirds [19]. The independent molecular dynamic simulation of ErCry4 mutants at residue 317 and 320 further suggests an important role of these two residues given their influence on fluctuation changes in the protein.

Mutational studies in chicken Cry4 suggested a reversible light dependent conformational change in the C-terminal region [58] and we speculate that the identified sites at the intrinsically disordered C-terminus (e.g. 510, 525) could very well be directly involved with the light-activated function of Cry4 and interaction partner binding since it was shown that the C-terminus of Cry4 interacts with a G-protein coupled receptor important for signaling [57,59].

Other aspects of our results also support a fundamentally different role of Cry4 compared to Cry1 and Cry2. Cry1 and Cry2 are present in all bird species suggesting that they serve at least one function essential for the survival of all birds as expected for clock proteins. In contrast, Cry4 was independently lost in at least three clades: hummingbirds, parrots and Tyranni. The clades that independently lost Cry4 are all characterized by long resident and tropical histories and may have lost the potential magnetoreceptor as the Earth’s magnetic field close to the magnetic equator is less useful for magnetic compass orientation than at more temperate latitudes [60,61]. However, other basal and resident tropical species including the cassowary, ostriches and tinamus have kept an intact and complete Cry4 and thus residency alone is unlikely to be the main driver behind Cry4 loss. It is expected that Cry4 most likely serves several different functions as it is expressed in all tissues investigated so far [22], and it is important to keep in mind that light dependent magnetic sensing is probably not a sensory capacity that is exclusive to migratory songbirds [30,62,63] but see [21] page S157). Both scenarios could explain conservation of Cry4 also for basal, resident species.

Regarding the Cry4-losses it is particularly interesting that, within the Tyranni, long-distance, night-migratory tyrant flycatchers recently evolved [40,41], which show very similar behaviors to night-migratory songbirds. If Cry4 is the magnetoreceptor in night-migratory songbirds, this recent appearance of a night-migratory life-style must have raised the necessity to either evolve magnetoreception-independent navigation or to evolve an independent magnetosensory system distinctly different from night-migratory songbirds. The loss in this group however shows, that Cry4 is not essential for night migration in passerines. Nevertheless, the protein’s major importance for night time magnetic compass orientation in songbirds is still supported, as songbirds evolved their migratory phenotype in the presence of Cry4. We will further investigate these finding experimentally in the field in the coming years to test if tyrant flycatchers are able to use the Earth’s magnetic field as a compass cue for orientation - and if so, whether their magnetic compass is a light-dependent [64] inclination compass [60] that can be disrupted by radiofrequency magnetic fields [65-69]. The tyrant flycatchers lacking Cry4 provide perfect natural knockouts of the putative magnetoreceptor Cry4 to test hypotheses of the radical-pair based magnetic compass mechanism, which has the potential to offer major support for Cry4 as a magnetoreceptor, and to provide a tool to study the evolutionary history of orientation and navigation mechanisms in birds.

## Methods

### Cryptochrome sequence extraction and alignment

We use the recently published dataset of 363 avian genomes assembled within the Vertebrate Genome Project (VGP/B10K) [32], (accession PRJNA545868) as the core dataset for our analyses. We extracted the coding sequence of all Cry members known to be present in birds, specifically Cry1a, Cry1b, Cry2, Cry4a and Cry4b. We used an in-house script based on blastn [70], which we optimised to extract Cry4a and b and validated nonsense mutations in the extra exon of Cry4b with whole genome resequencing data of species with high quality genomes and resequencing data available (see supplementary methods for details). As a query, we used the respective Cry sequences of the European robin (*Erithacus rubecula*, accession number PRJEB38659). Codon alignments were generated with mafft [71] and the MUSCLE algorithm in MEGA followed by polishing by hand including the exclusion of too short sequences (i.e. just one short exon present). To verify losses of Cry4 in single clades we used high quality reference genomes from the VGP genomeark website (https://vgp.github.io/genomeark/). To further investigate whether Cry4 was lost via gene deletion or pseudogenization in Tyranni we generated a synteny of two high quality reference genomes that had no sequencing gaps in the Cry4 area with Satsuma [72], including one species with Cry4 and one species that had lost it. We used the ConSurv server to quantify and visualize variability and differences between the Cry alignments [73].

A consensus species tree with 50% majority rule was generated based on 100 trees received from birdtree.org based on the Ericson all species data set with a set of 10000 trees and 9993 OUTs each [74]. Branch lengths for the constrained species tree were calculated based on the Cry4 nucleotide and amino acid alignment with iqtree, using the best-fitting nucleotide and amino acid substitution model identified by the program (GTR+F+R10 and JTT+R6). These branch lengths were necessary to test for evolutionary rate shifts.

### Characterization of migratory phenotype

Bird species were classified according to their migratory behaviour: Obligate migratory and obligate resident species were classified accordingly, partial migrants with most of the species being migratory were classified as migratory. Species for which a distinct classification is difficult (e.g. nomadic and irruptive species) were classified as ‘indistinct’.

### Molecular evolutionary analysis

#### Ancestral state reconstruction

For visualization and as input for some programs we reconstructed the ancestral migratory phenotype (migratory/resident) and Cry4a presence/absence. through the maximum likelihood program ace from the R-package “phytools” [75,76] with a specifically designed Q-matrix (transition matrix, explaining how traits evolve back and forth, see supplementary methods for more detail). For reconstruction of single amino acid sites, we used the output of codeml [46].

#### Positive selection

We tested for positive selection by fitting codon substitution models in the codeml package of PAML [45,46]. Omega is defined as the nonsynonymous/synonymous rate ratio (omega = dN/dS) and is an important indicator of the type of selective pressure at the protein level, with omega = 1 meaning neutral evolution, omega < 1 purifying selection, and omega > 1 diversifying positive selection. We used two different models: 1) The site-model allows for each codon to evolve under different omega values shared by all branches, and 2) the branch-site model which can detect different selection regimes in particular lineages on foreground branches, i.e. the branches of interest in the tree. Foreground branches are the selected branches of interest or particular lineage(s), all other branches are referred to as background branches. Background branches share the same distribution of omega (dN/dS) value among sites, whereas different values can apply to the foreground branch.

For the site-model, the nested models M1a (omega <=1) and M2a (omega > 1 allowed) were run and the negative log-likelihood was compared with a likelihood ratio test (LRT). P-values were drawn from a χ^2^ distribution and the model that significantly fitted the data better was selected [77]. If the LRT allows for omega > 1, a Bayes Empirical Bayes (BEB) analysis was used to compute the posterior probability for each site to be in this class (omega >1). Also, the more parameter-rich nested site-models M7 and M8 were compared in the same way for Cry4. The site-models M1a and M2a were applied to Cry4a, Cry1a and b and Cry2 [46] (see Tables S1-S3 for details).

For the branch-site model we chose specific foreground branches for which positive selection is allowed in the more complex model, while omega is restricted to <=1 on background branches. As foreground branches we tested (i) all passerines, (ii) clades with mostly migratory species within the passerines, (iii) the branch leading to the passerines, and (iv) branches to clades with mostly migratory species within the passerines. We additionally selected (v) the non-passerines as foreground clades and tested this against all passerines. For all runs, two nested models were tested and the log-likelihood was compared as described above [77]. The branch-site model was run only for Cry4a, as the site-models already struggled to converge for Cry1a, Cry1b and Cry2 due to extremely high sequence conservation (Table S2).

We also used the Hyphy [47] FEL and Contrast-FEL package to test for positive selection on passerine foreground branches compared to non-passerines, and to test for sites that changed selection pressure between the two groups (see supplementary methods and results, Tables S4-S5).

#### Evolutionary rate shift

Another approach that allows us to characterize functional changes between groups detects shifts in the evolutionary rate. We applied the RateShift method [78] on amino-acid sequence alignments, implemented using the Bio++ libraries (https://github.com/BioPP/rateshift). The rate shift is detected between focal foreground and background branches. We conducted three independent analyses, to test whether the rate of amino-acid substitutions changed on the branches including the passerines clade, (ii) clades with a passerine migratory ancestor, and (iii) only passerine night migrants. Due to the lack of sequence variability of Cry1 and 2 revealed in the steps above, this model was applied only to Cry4.

#### Correlation of Cry4b extra exon loss and migratory phenotype

To test for correlated evolution of migratory phenotype and extra exon of Cry4b present or absent, we first tested the phylogenetic constraints of the two traits through the D-statistic implemented in the phylo.d package [44] using 1000 permutations. We used the maximum likelihood method of BayesTraits [75,79] to test for the correlation of the two discrete traits, taking the phylogenetic relationship into account: Cry4b extra exon loss (either by deletion or accumulation of nonsense mutations) and migratory phenotype. We included only songbird (Oscine) species that have a clear migratory or resident phenotype (n = 128). To prevent the transitions from an absent or non-functional extra exon to a present exon, the parameters explaining these two transitions were restricted to 0.00. To test for dependence of trait evolution, a model of independent trait evolution was compared to a model with dependent trait evolution via a likelihood ratio test. The p-value was drawn from a χ^2^ distribution and the model that significantly fitted the data better was chosen.

#### Prove absence of Cry4 in tyrant flycatcher

We isolated RNA from brain, ovary and retina of a least flycatcher (Empidonax minimus) and sequenced libraries on a NovaSeq (Illumina) generating over 125M 2×150 PE reads per sample. We made a reference free transcriptome alignment with Trinity [80-83] and quality control with BUSCO and Bowtie [80-82]. Blastn was used to identify transcripts of Cry1a/b, Cry2 and Cry4 (see supplementary methods for more information).

#### Protein molecular dynamics simulation

While phylogenetic tools are very powerful and useful in investigating the evolutionary pressure on sequence differences on species scale, they fall short in revealing dynamic features of the proteins or how these are altered by single mutations. We thus complement our phylogenetic approach with molecular dynamics (MD) simulations, that provide an efficient way of investigating protein models employing the NAMD package [83,84] through the VIKING interface [85] to investigate the dynamics of Cry4. Using the already crystalized pigeon Cry4 as template [27], we apply homology modeling to create a high-quality European robin Cry4 model based on its amino acid sequence [19,26,48] and investigate changes in the fluctuation at each site after artificial mutations at identified relevant sites are introduced (see supplementary methods and results for more information and model evaluation).

## Acknowledgments

This study would not have been possible without the fantastic resources of 363 genomes, released by the B10K Consortium in 2020. We are extremely grateful for the encouragement, support and discussions we have received from the consortium throughout the project; special thanks go to Shaohong Feng and Qi Fang for running accompanying analysis on synteny. We thank Benjamin Van Doren and Bronwyn Butcher, who aided organization of tissue samples for RNAseq experiments, the Ornithology Collection of the Cornell University Museum of Vertebrates for sample provision, and Cornell’s TREx facility for sample processing and sequencing. Thanks also go to Barbara Helm, Arne Hegemann and Jan von Rönn for feedback and discussion on species specific classification of migratory behavior. We thank the Max Planck Society (MPRG grant MFFALIMN0001 to ML), the Volkswagen Foundation (Lichtenberg professorship awarded to IAS), the Deutsche Forschungsgemeinschaft (GRK1885 Molecular Basis of Sensory Biology awarded to HM and IAS; and SFB 1372 Magnetoreception and Navigation in Vertebrates, no. 395940726 awarded to HM, ML and IAS), the Ministry for Science and Culture of Lower Saxony (Simulations meet experiments on the nanoscale: opening up the quantum world to artificial intelligence (SMART) awarded to IAS), and the European Research Council (under the European Union’s Horizon 2020 research and innovation program, grant agreement no. 810002, Synergy Grant: ‘QuantumBirds’, awarded to H.M.). Computational resources for the simulations were provided by the CARL Cluster at the Carl-von-Ossietzky University, Oldenburg, supported by the DFG and the Ministry for Science and Culture of Lower Saxony. This work was also supported by the North-German Supercomputing Alliance (HLRN).

