## Supplementary Information for "Evidence for adaptive evolution towards high magnetic sensitivity of potential magnetoreceptor in songbirds"

**This PDF file includes:**

Supporting text
Figures S1 to S6
Tables S1-S6
SI References

**Other supporting materials for this manuscript include the following:**

Datasets S1 to S2
Alignment S1 to S4

**Cryptochrome extraction and alignment** Here, we leverage the recently curated dataset of 363 avian genomes assembled within the Vertebrate Genome Project (VGP/B10K) (Feng et al., 2020, accession PRJNA545868). We used this dataset to extract full genomic coding sequences for all members of the cryptochrome family, Cry1a and b, Cry2 and Cry4a and b that have been described in birds using the sequence from the European robin *Erithacus rubecula* as queries, a night-migratory songbird that has been focally studied in the context of magnetic compass orientation (European Nucleotide Archive: accession number PRJEB38659).

We used blastn (Zhang et al, 2000) for sequence extraction and optimized parameters for Cry4a. Specifically, we compared Cry4a mRNA data from the NCBI database to the blastn search of the robin Cry4a as query against the genomes of the corresponding species, and to recover the whole sequence the blastn option -word_size was set to 10. The resulting output was filtered for the best fitting hits, using the e-value and percent identity. Additionally, the positions of Cry4a hits were compared to Cry1 (a and b) and Cry2 hits, to exclude the possibility of accidentally mistaking Cry1 or Cry2 exons for Cry4, and the correct order of exons was checked based on the position of blast hits on the scaffolds. We used the same script to receive Cry1 and Cry2 sequences. AUGUSTUS, a gene-prediction program (Stanke et al, 2004), was run specifically on the part of the genome where Cry4a hits were identified by blastn to find additional exons that were too different for blastn to find, and to identify start and stop codons. To completely isolate the extra exon of Cry4b the script was adapted to also find exons with larger gaps, as the exon was highly diverse and fragmented for many species. To verify losses of Cry4 in single clades, high-quality reference genomes of representative species belonging to these clades from the VGP genomeark website (https://vgp.github.io/genomeark/) were processed in the same manner as above. To investigate whether Cry4 was lost via a gene deletion in basal passerines a synteny of high-quality reference genomes was generated with Satsuma (Grabherr et al., 2010), including one species with Cry4 and one species that had lost it. Also, the presence of ten adjacent genes on the same scaffold was investigated for both species. To confirm deleterious mutations (e.g. frameshift or nonsense mutations) in the extra exon of Cry4b, IsoSeq, RNASeq and whole genome resequencing data of the Eurasian blackcap (Delmore et al., 2020a), zebra finch (Singhal et al., 2015) and European robin (not published) were analyzed. We used the CunSurv server to quantify and visualize differences between the Cry alignments (Ashkenazy et al., 2016).

Based on the extracted exons for all cryptochrome genes in focus, codon alignments were generated with mafft (Katho et al., 2002). The codon alignment was carried out using the MUSCLE codon aligner of MEGA, and a subsequent polishing by hand in MEGA, including the exclusion of too short sequences (i.e. just one exon present). For tests of selection and trait mapping, a consensus species tree with 50 % majority rule was generated based on 100 trees received from birdtree.org (Jetz et al., 2012) based on the Ericson all species data set with a set of 10000 trees and 9993 OUTs each. The selection analysis in PAML was also carried out using the species tree generated by the B10K (Feng et al., 2020) to validate the robustness of the results. Branch lengths for the first constrained species tree were calculated based on the Cry4 nucleotide and amino acid alignment with iqtree (Nguyen et al., 2015), using the best-fitting nucleotide and amino acid substitution model identified by the program (GTR+F+R10 and JTT+R6).

We validated the loss of Cry4 using high-quality reference genomes of representative species for all three groups: the Annas hummingbird *Calypte anna* (Trochilidae, GCA_003957555.2), budgerigar *Melopsittacus undulatus* (Psittaciformes, GCA_012275265.1), and lance-tailed manakin *Chiroxiphia lanceolata* (Suboscines, GCA_009829145.1), respectively.

### Molecular evolutionary analysis

**Ancestral state reconstruction** A phylogenetic trait reconstruction with the maximum likelihood ace program from the R-package “phytools” (Pagel, M., 1994) was carried out reconstructing the migratory phenotype and the state of Cry4a (absent, present, fragmented), with both phylogenies. The B10K phylogeny has the advantage, that branch length represent species and not protein divergence which is preferred for trait reconstructions. For the reconstruction of the Cry4a state, a transition matrix was designed to allow only changes from present to fragmented and absent, and from fragmented to absent but not vice versa as it is not possible to regain the protein once it was lost. The migratory phenotype was mapped onto the species tree and the maximum likely node states were visually compared between the two phylogenies. As the reconstruction of traits was robust between phylogenies, the displayed results refer to the B10K phylogeny. For reconstructions of single amino acid sites, we used the output of codeml. Ancestral codon states are inferred by a likelihood/empirical Bayes approach in PAML based on a codon model (codon frequency model F3x4) and a discrete gamma distribution for rate variations among sites (Yang, 1997, 2007).

### Positive selection with Hyphy FEL and Contrast-FEL

For additional evidence, we also used FEL from HyPhy (Pond and Muse, 2005) to test for selection on foreground branches of passerines compared to non-passerines, similar to the branch-site model. We also used Contrast-FEL to test for sites under divergent selection pressure between the two groups.

The results of FEL were concordant with the branch-site model (table S4) and sites 320, 330, 425, 510 and 512 were under positive selection in the passerine clade. The Contrast-FEL analysis identified 11 sites with a significant shift of selection pressure between non-passerines and passerines (with q-value < 0.02 to control for false positives, table S5). This includes site 320, concordant with the shift from negative to positive selection (branch-site test PAML), 510 and 525.

**Directional selection** Migratory behavior likely evolved several times independently in birds, which can be considered as convergent evolution with directional selection for the optimal amino acids. Directional selection can be characterized by only few substitutions in the phylogeny, evolving towards the most optimal amino acid state given a certain phenotype or environmental condition.

Models that aim for detecting this pattern link a trait of the species to a change of fitness at any given amino acid site. The analysis was conducted with the program coevol, which is a Bayesian inference program that uses Markov Chain Monte Carlo methods (Parto and Lartillot, 2018). All passerines that show a clear resident (93 species) or migratory (31 species) phenotype were included in the analysis and ancestral behavior (migratory/resident) was reconstructed using the same methods as described above. Our resulting tree was labelled accordingly and used as input for the differential selection diffsel model of the program coevol. The Differential Selection model was used with three groups (DS3) as recommended by Parto and Lartillot (2018). These three groups were split into the interior branches to calculate the baseline condition, as well as resident and migratory species. 15000 iterations were done and the program tracecomb was used with a burn-in of 3000 (20 % of iteration number) to check whether the chain had converged well. This model was applied to Cry4a, Cry1a, Cry1b and Cry2.
We identified four sites (residues 116, 202, 516 and 525) that are under directional selection in Cry4a and which have a higher fitness factor in migratory songbirds (posterior probability (PP) $> 0.95$ or $< 0.05$ in all six runs, Table S1). For Cry1a, Cry1b and Cry2 we did not identify any sites (with PP $> 0.95$ or $< 0.05$) in this context.

**Co-evolution of candidate sites** To test whether our identified candidate sites under positive selection co-evolved, we used the program comap. This approach applies probabilistic substitution mapping to detect patterns of co-substitutions (Dutheil et al., 2005; Dutheil and Galtier, 2007) and we applied the clustering and candidate-based approach. The clustering method does not require previously defined groups of candidate sites. For the candidate-based approach, we selected the four symmetrical sites that we identified as being under positive selection (entry ID: 68, 83, 116, 189, see results and figure 4). For both approaches, we tested for correlative co-evolution, which assumes that sites tend to undergo changes in the same branches of the tree, and compensating mutations, which accounts for the chemical properties (polarity) of the amino acids. We conducted 10000 simulations.

**Prove absence of Cry4 in tyrant flycatcher**

**RNA isolation and library preparation and sequencing** RNA was purified from brain, retina and ovary tissue of a female least flycatcher museum specimen from the collection of the Cornell Museum of Vertebrates (specimen CUMV 55184) preserved in RNAlater using Trizol (Thermo Fisher) according to the commercial protocol with the following additions: after the first phase separation, an additional chloroform extraction step of the aqueous layer in Phase-lock Gel heavy tubes (Quanta Biosciences); an addition of 1ul Glyco-blue (Thermo Fisher) immediately prior to isopropanol precipitation; two washes of the RNA pellet with 75% ethanol. Quality control was confirmed with spectrophotometry (Nanodrop) and a Fragment Analyzer (Agilent).

Tru-Seq-barcoded RNAseq libraries were generated with the NEBNext Ultra II Directional RNA Library Prep Kit (New England Biolabs) with polyA enrichment using 1ug total RNA as input. Each library was quantified with a Qubit 2.0 (dsDNA HS kit, Thermo Fisher) and the size distribution was determined with a Fragment Analyzer prior to pooling. The libraries were sequenced on a NovaSeq (llumina) generating over 125M 2x150 PE reads per sample.

**Least flycatcher transcriptome assembly** The following analyses were conducted on all three tissues separately and all data combined respectively. To assess quality, we conducted fastgc for each tissue. A de novo, reference free assembly of the transcriptome was created with trinity (Grabherr et al., 2011) using the –trimmomatic option to trim the reads (e.g. remove adapter sequences). The assembly quality was assessed by remapping the reads to the assembly with Bowtie (Langmead et al., 2009). We ran BUSCO on the final assembly to check for completeness (Manni et al., 2021), and used blastn to identify transcripts of Cry1a/b, Cry2 and Cry4.

### Molecular dynamics study of ErCry4

The atomistic structure of the European robin cryptochrome 4 (ErCry4) was constructed computationally from the known primary amino acid sequence (Günther et al. 2018). A homology modelling approach was used to reconstruct the ErCry4 structure, where the recently crystalized pigeon cryptochrome 4 (Zoltowski et al. 2019) was used as the molecular template; the obtained wild type ErCry4 structure was thus equivalent to the structure simulated and discussed in a recent publication (Günther et al, 2018; Xu et al. 2021; Hanić et al., 2022). The ErCry4 structure was solvated and neutralized in a water box of 92 Å × 105 Å × 100 Å by using 0.15 M of NaCl. The system was then treated dynamically to ensure its stability by using the program NAMD 2.10 for the wildtype simulations and 2.13 for the mutations (Phillips et al. 2005). The initial structure minimization and equilibration protocols were largely adapted from earlier studies (Nielsen et al. 2018, Frahs et al. 2019, Korol et al. 2019, Frederiksen and Solov'yov 2020, Xu et al. 2021). The simulations that were used for data sampling (production simulation) were carried out for 200 ns in the NVT statistical ensemble using a Langevin piston to ensure a constant temperature of 310 K (Nosé et al. 1983). The mutation introduced was made from the last frame of the 200 ns production run and it was introduced using the VMD mutator tool (Humphrey et al. 1996). The mutated simulation was carried out using the same simulation scheme as for the wildtype ErCry4. For a more complete overview of the simulations see Table S6. Particle mesh Ewald was used in all simulations to treat long range electrostatics with a cut-off distance of 12 Å (Darden et al. 1993). CHARMM36m force fields were used to describe the atoms in the protein and water molecules (Denning et al. 2011, Hart et al. 2012, Huang et al. 2017). The force field parameters for the FAD cofactor taken from earlier successful studies (Solov'yov et al. 2012, Nielsen et al. 2017, Kattnig et al. 2018, Nielsen et al. 2018, Günther et al. 2019). The analysis of planarity of the four investigated amino acid residues (Figure. 4) was done using in-house scripts, the stability was calculated using the RMSD tool in VMD (Humphrey et al. 1996) and the RMSF was calculated using the ProDy python package (Bakan, et al. 2014).

### Computational results

The stability of the protein structures was estimated through the time evolution of the root mean square deviation (RMSD) of the structures. The RMSD was calculated using VMD (Humphrey et al. 1996) as follows:

$$\mathrm{RMSD} = \sqrt{\frac{1}{N}\left(\sum_{i=1}^{N} \vec{r}_{\mathrm{i}}(t) - \vec{r}_{\mathrm{i}}(t=0)\right)^{2}}\ , \qquad (1)$$

where N is the total number of backbone atoms in a protein, $\vec{r}_{\mathrm{i}}(t)$ is the vector describing the coordinates of the i'th atom in the backbone. Figure S5 shows the calculated time evolution of the RMSD for the ErCry4 and ClCry4 wildtype (WT) cryptochromes as well as the ErCry4 C317H, E320K and ClCry4 H317C mutations.

Figure S5 reveals that all studied structures are stable as no noticeable increase of the studied dependencies was observed. It is noticeable that the time evolution for the ClCry4 structure behaves similarly to the other proteins, while ClCry4 is the only avian cryptochrome for which a crystal structure is available. Following the RMSD calculation, the root mean square fluctuations (RMSF) of the structures was calculated as (Bakan et al. 2011, Bakan et al. 2014, Zhang et al. 2021):

$$\mathrm{RMSF(i)} = \sqrt{\frac{1}{T}\sum_{t_j=1}^{T} \left|\vec{r}_{\mathrm{i}}(t_j) - \overrightarrow{r_{\mathrm{i}}^{ref}}\right|^{2}}\ , \qquad (2)$$

here T is the total time of the trajectory $\overrightarrow{r_{\mathrm{i}}^{ref}}$ is the reference position of particle i and $\vec{r}_{\mathrm{i}}(t_j)$ is the position of particle i after a time $t_j$. The RMSF has been calculated for 20 different subsampling schemes, by splitting each trajectory into 20 equal-sized pieces and calculating the RMSF for each of them, the final plot is thus found by averaging over the 20 points giving an average (bold lines) and a standard deviation from the average (transparent area around graphs).

The RMSF revealed several interesting features for the structures analyzed. It is observed that the ErCry4 C317H mutation did not stabilize the same parts as the ClCry4 H317C mutation, The ErCry4 C317H mutation saw an extinction of two

RMSF peaks (residues 189 and 202) whereas the ClCry4 H317C mutation saw a stabilization in the region around residue 230, also known as the phosphate binding loop (Schuhmann et al., 2021; Hanić et al., 2022a,b). Furthermore, the ErCry4 K320E mutation revealed that even though two mutation sites are very close to each other they will not have the same impact on the flexibility of the structure as the ErCry4 K320E mutation only saw stabilization of the 202 region while the 189 region remained flexible. Comparing the two ErCry4 mutation to the ClCry4 H317C mutation it is observed that the ClCry4 mutation did not see any significant stabilization at the 189 site but saw a similar stabilization at the 202 site. Furthermore, it is observed that the ClCry4 H317C mutation stabilizes around residue 89 which neither of the ErCry4 mutations did. The planarity of the four sites (68, 83, 116 and 189) in ErCry4 was also investigated as it turned out striking that these residues seem to follow a single plane. The planarity was investigated by analyzing the dynamics of the protein and placing two planes passing through the centres of masses of residues 68, 83, 116 and 83, 116 and 189. The angle $\alpha$ between the two planes determines the deviation of the four residues from planarity and is calculated as

$$\alpha = \mathrm{acos}(\overrightarrow{n_1} \cdot \overrightarrow{n_2})\,, \tag{3}$$

where $\overrightarrow{n_1}$ and $\overrightarrow{n_2}$ are the normalized normal vectors for the two planes introduced above. Figure S6 (right) shows the time evolution of the angle between the two planes described by Eq. (3).

What is important to note from Figure S6 (right) is not only the fact that the angle is indeed stable with it being less than 5 degrees for most of the trajectory. Another way to check the planarity of the four residues is to, once again, calculate a plane using the plane spanned by residues 68, 83 and 116 and calculating the shortest distance from the fourth residue to said plane. The plot as well as distribution of the shortest distance can be seen on Figure S6, again it is confirmed that the residues are mostly planar.

$$d = \vec{n} \cdot \vec{v} = X_n \cdot x + Y_n \cdot y + Z_n \cdot z, \tag{4}$$

where d is the distance, $\vec{n} = \begin{pmatrix} X_n \\ Y_n \\ Z_n \end{pmatrix}$ is the normal vector of the plane spanned by residues 68, 83 and 116 and $\vec{v} = \begin{pmatrix} x \\ y \\ z \end{pmatrix}$ is the vector spanning from residue 189 to the aforementioned plane. The inner product between the vector $\vec{v}$ and the normal vector of the plane gives the length of the projection of the vector, $\vec{v}$, on to the normal vector $\vec{n}$ of the plane, which by definition is the shortest distance given in Eq. (4). From Figures S6 it is evident that some kind of planarity exists for the four residues, and that after some equilibration time this planarity is quite stable although what the planarity means for the overall protein dynamics is a yet unexplored area.

**Dataset S1 (separate file).** Table with cryptochrome sequence extraction information

**Dataset S2 (separate file).** Results of RateShift comparison of passerines vs non-passerines

**Alignment S1 (separate file).** Cry4 alignment

**Alignment S2 (separate file).** Cry1a alignment

**Alignment S3 (separate file).** Cry1a alignment

**Alignment S4 (separate file).** Cry2 alignment

**References only appearing in SI**


1. Feng, S., Stiller, J., Deng, Y., Armstrong, J., Fang, Q., Reeve, A. H., ... & Zhang, G. (2020). Dense sampling of bird diversity increases power of comparative genomics. Nature, 587(7833), 252-257.
2. Zhang, Z., Schwartz, S., Wagner, L., & Miller, W. (2000). A greedy algorithm for aligning DNA sequences. Journal of Computational biology, 7(1-2), 203-214.
3. Stanke, M., Steinkamp, R., Waack, S., & Morgenstern, B. (2004). AUGUSTUS: a web server for gene finding in eukaryotes. Nucleic acids research, 32(suppl_2), W309-W312.
4. Grabherr, M. G., Russell, P., Meyer, M., Mauceli, E., Alföldi, J., Di Palma, F., & Lindblad-Toh, K. (2010). Genome-wide synteny through highly sensitive sequence alignment: Satsuma. Bioinformatics, 26(9), 1145-1151.
5. Delmore, K., Illera, J. C., Pérez-Tris, J., Segelbacher, G., Ramos, J. S. L., Durieux, G., ... & Liedvogel, M. (2020a). The evolutionary history and genomics of European blackcap migration. Elife, 9, e54462.
6. Singhal, S., Leffler, E. M., Sannareddy, K., Turner, I., Venn, O., Hooper, D. M., ... & Przeworski, M. (2015). Stable recombination hotspots in birds. Science, 350(6263), 928-932.
7. Ashkenazy, H., Abadi, S., Martz, E., Chay, O., Mayrose, I., Pupko, T., & Ben-Tal, N. (2016). ConSurf 2016: an improved methodology to estimate and visualize evolutionary conservation in macromolecules. Nucleic acids research, 44(W1), W344-W350.
8. Katoh, K., Misawa, K., Kuma, K. I., & Miyata, T. (2002). MAFFT: a novel method for rapid multiple sequence alignment based on fast Fourier transform. Nucleic acids research, 30(14), 3059-3066.
9. Jetz, W., Thomas, G. H., Joy, J. B., Hartmann, K., & Mooers, A. O. (2012). The global diversity of birds in space and time. Nature, 491(7424), 444-448.
10. Nguyen, L. T., Schmidt, H. A., Von Haeseler, A., & Minh, B. Q. (2015). IQ-TREE: a fast and effective stochastic algorithm for estimating maximum-likelihood phylogenies. Molecular biology and evolution, 32(1), 268-274.
11. Pagel, M. (1994). Detecting correlated evolution on phylogenies: a general method for the comparative analysis of discrete characters. Proceedings of the Royal Society of London. Series B: Biological Sciences, 255(1342), 37-45.
12. Yang, Z. (1997). PAML: a program package for phylogenetic analysis by maximum likelihood. Computer applications in the biosciences, 13(5), 555-556.
13. Yang, Z. (2007). PAML 4: phylogenetic analysis by maximum likelihood. Molecular biology and evolution, 24(8), 1586-1591.

### Supporting figures with references in main manuscript

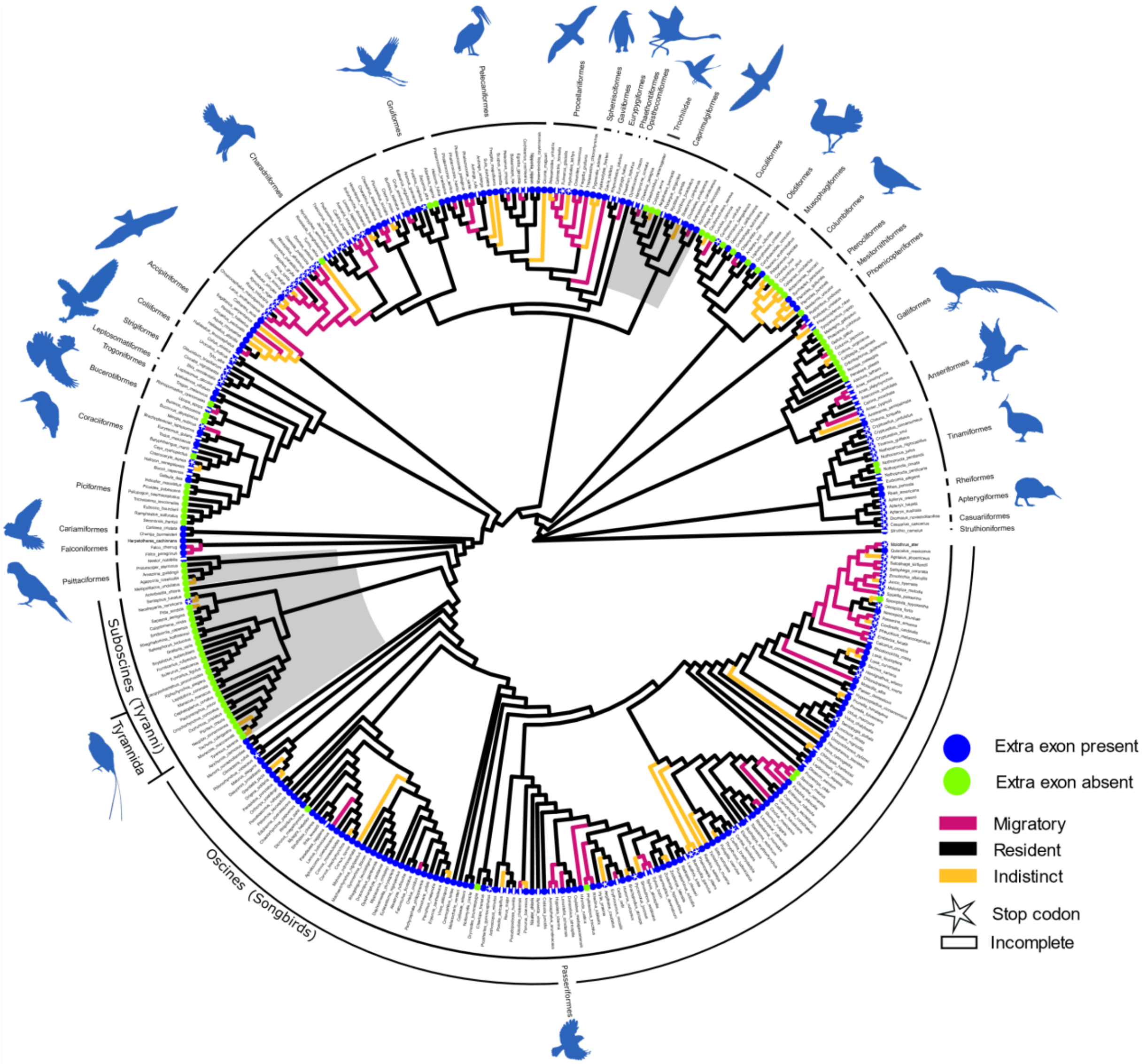


*Figure S1:* ***Loss of the additional exon found in Cry4b.*** *The extra exon indicative for Cry4b is highly variable in all species (exon present = blue) and its evolution is characterized by multiple independent losses (green) and by accumulation of nonsense mutations (indicated by star = stop codon and bar = deletions).*

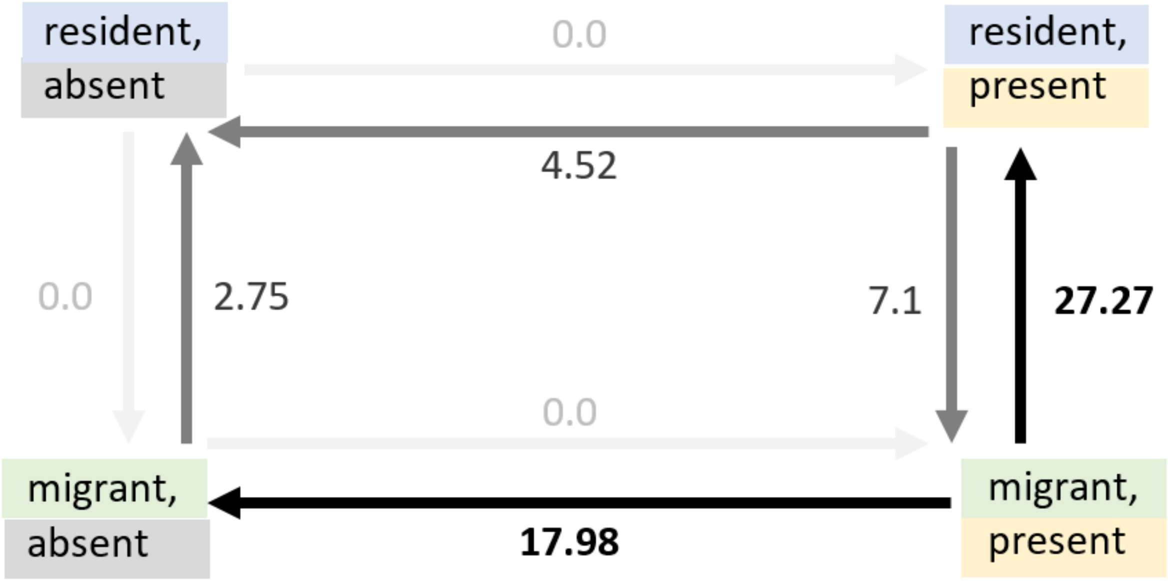


*Figure S2:* ***The loss of the extra exon in Cry4b is significantly correlated with migratory behavior in songbirds (Oscines).*** *The numbers at the arrows indicate transition rates between being migratory or resident and the presence or absence of Cry4b. If Cry4b is present in a migrant, we observe a high transition likelihood from being migratory to being a resident (rate 27.27, dimensionless). Furthermore, if you are a migrant with Cry4b present there is a high likelihood that you would lose Cry4b (rate 17.98). This could suggest an evolutionary disadvantage and selection against having Cry4b if you are a migrant. Transitions from a lost extra exon to a present extra exon were restricted to 0. LRT: 15.37, df: 3, p-value: 0.0015.*

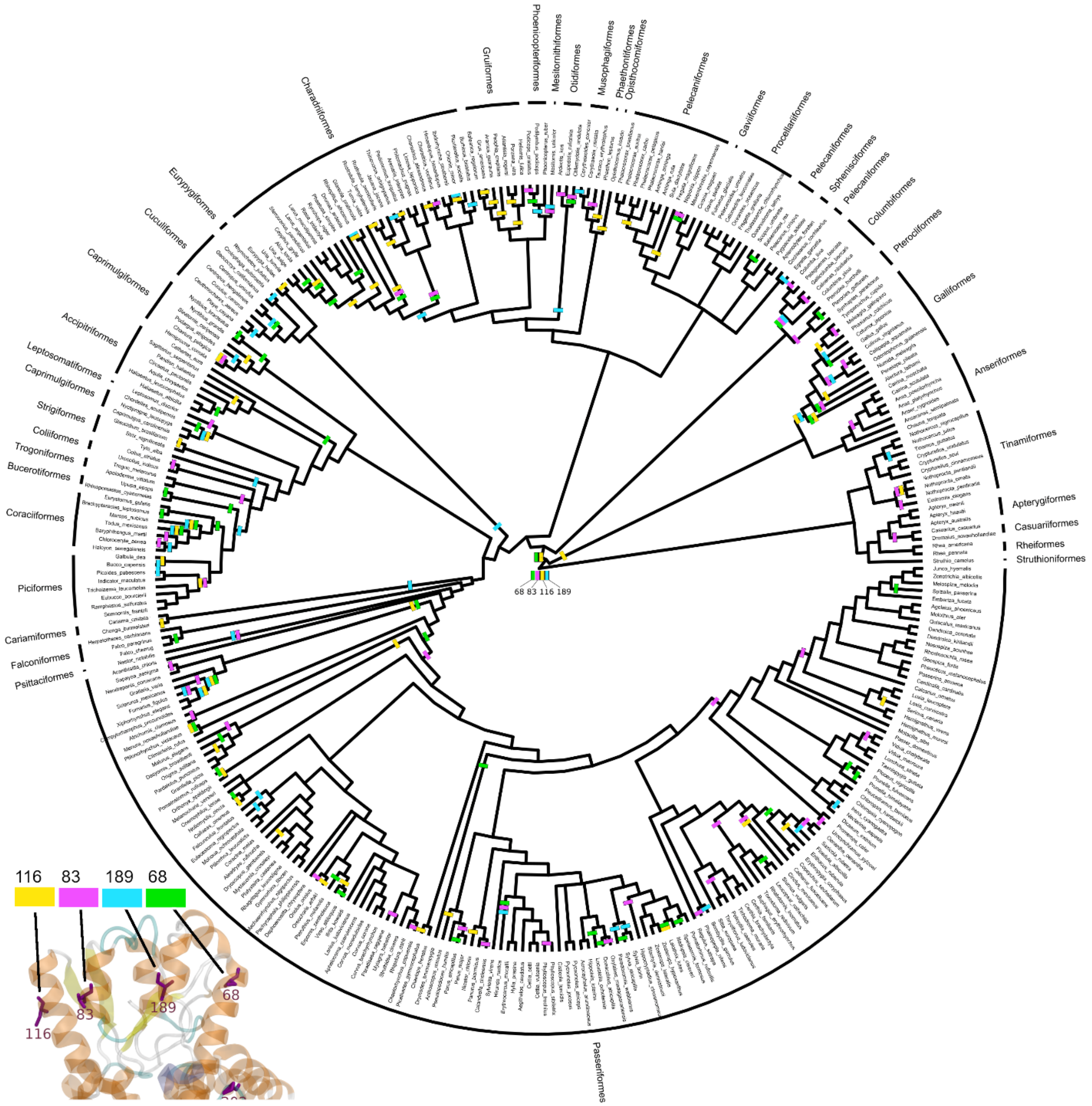


*Figure S3:* ***Do the substitutions of the four sites under positive selection that are arranged in a planar pattern (68- green, 83 - purple, 116 - yellow, 189 - blue) show signatures of convergent evolution (i.e. do they change together on the same branch)?*** *A bar (the color identifies the site at which an aa changed) on a branch in the phylogeny indicates that one substitution among the four planar arranged residues occurred. For example, on the first branch leading to Galliformes a substitution at site 116 (yellow) occurred whereas on the first branch to all other clades two substitutions happened at site 116 and 69. If the four sites coevolved, we would expect simultaneous substitutions of the four sites on the same branches. However, we could not detect such co-evolutionary patterns.*

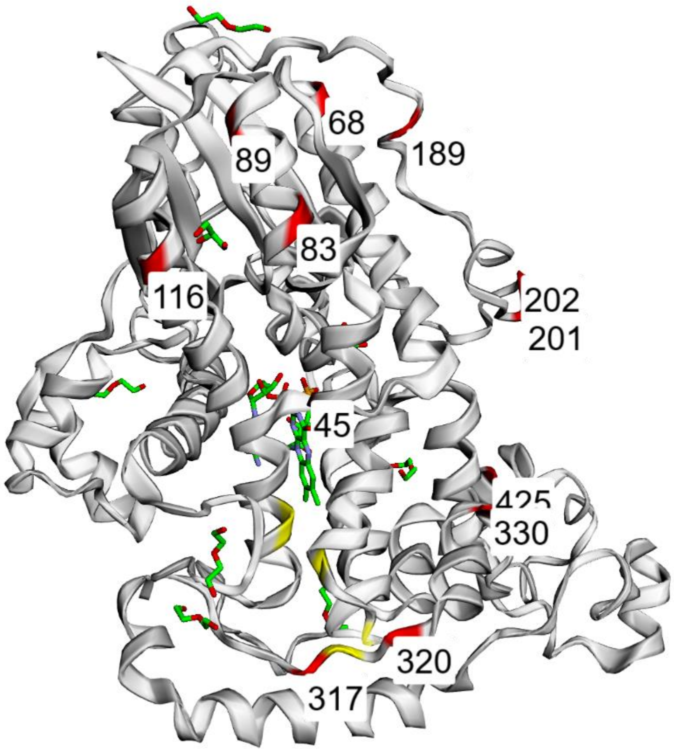


*Figure S4:* ***All candidate sites identified in Cry4 listed in table S1****. The candidate sites are highlighted in red. The tryptophan tetrad is colored in yellow.*

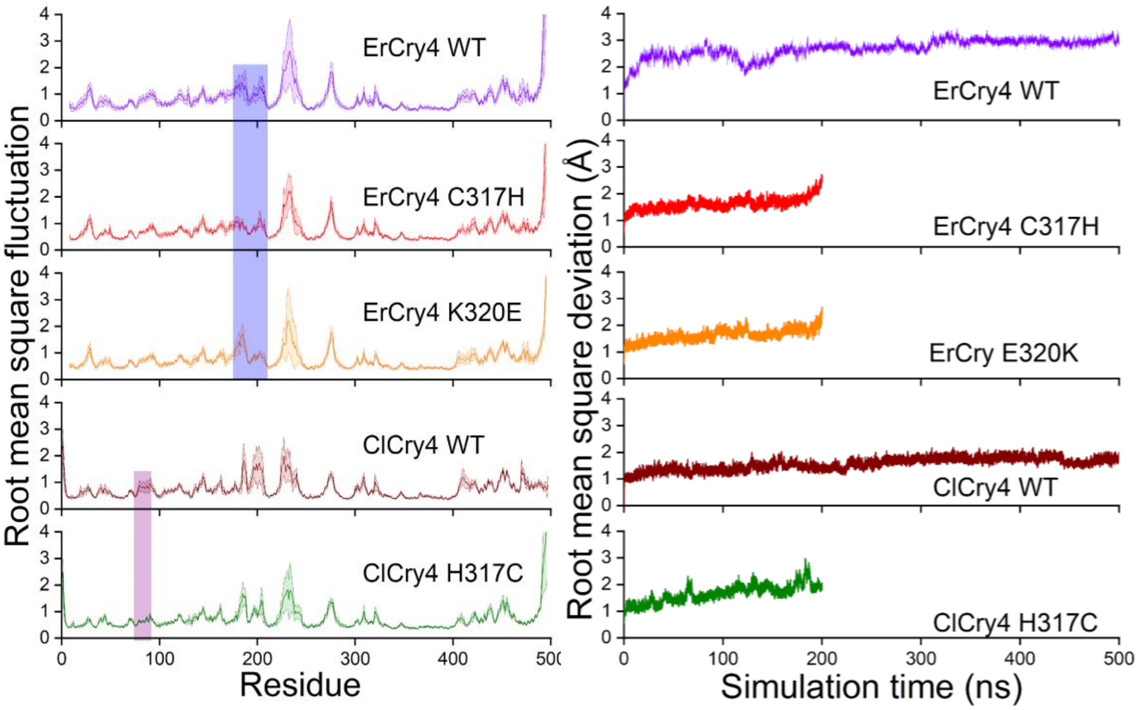


*Figure S5:* ***Calculated time evolution of the RMSF for the ErCry4 and ClCry4 wildtype (WT) and mutants.*** *Left: Root mean square fluctuations (RMSF) for the residues from ErCry4 and ClCry4 wildtypes as well as their mutations considered in this study. The RMSF has been calculated by means of Eq. (2). The RMSF revealed several interesting features for the structures analyzed. It is observed that the ErCry4 C317H mutation did not stabilize the same parts as the ClCry4 H317C mutation. The ErCry4 C317H mutation saw an extinction of two RMSF peaks (residues 189 and 202) whereas the ClCry4 H317C mutation saw a stabilization in the region around residue 230, also known as the phosphate binding loop. Furthermore, the ErCry4 K320E mutation revealed that even though two mutation sites are very close to each other they will not have the same impact on the flexibility of the structure as the ErCry4 K320E mutation only saw stabilization of the 202 region while the 189 region remained flexible. Comparing the two ErCry4 mutation to the ClCry4 H317C mutation it is observed that the ClCry4 mutation did not see any significant stabilization at the 189 site but saw a similar stabilization at the 202 site. Furthermore, it is observed that the ClCry4 H317C mutation stabilizes around residue 89 which neither of the ErCry4 mutations did. Right: Time evolution of the root mean square deviations (RMSD) computed for the crystal structure of wildtype pigeon cryptochrome (ClCry4), the ClCry4 H317C mutation, the wildtype European robin cryptochrome (ErCry4) and the C317H and E320H mutations of the ErCry4. The time evolution was performed over 200 ns for the aligned structure of the backbone of the protein structure. The RMSD has been calculated using Eq. (1).*

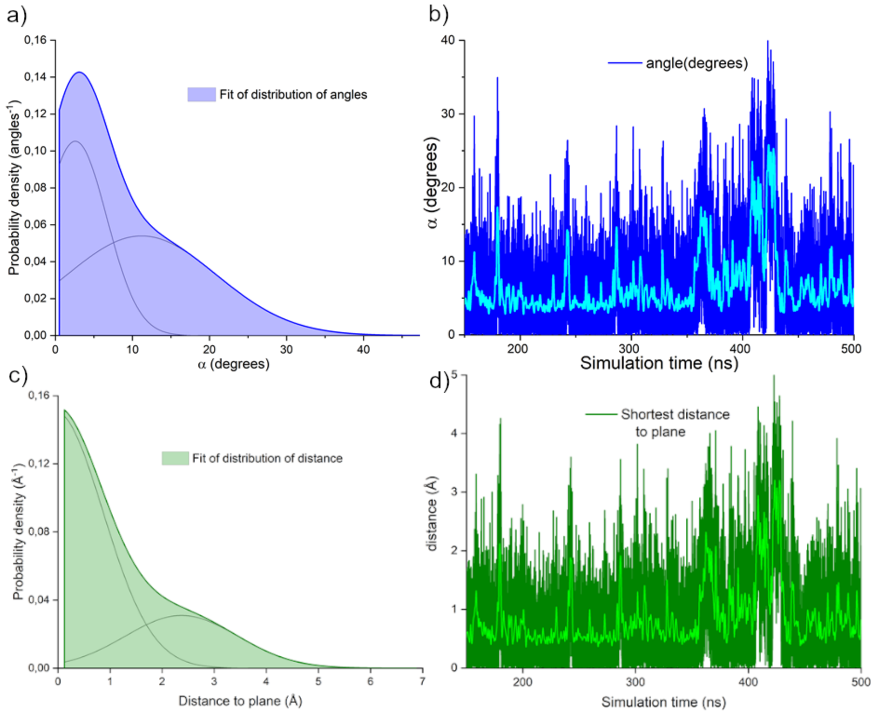


*Figure S6:* ***Planar pattern of four residues under positive selection.*** *a) Probability distribution of the angle α between the two planes spanned by residues 68, 83, and 116 and 68, 83 and 189, respectively. The curve is a sum of two gaussian distributions each shown in grey with the cumulative curve shown in blue and corresponds to the distribution of the angle in the figure to the right. b) Time evolution of the angle α between the two planes spanned by the four residues 68, 83, 116 and 189 in ErCry4. The first 140 ns of the trajectory has been omitted. Not only the fact that the angle is indeed stable with it being less than 5 degrees for most of the trajectory is important to note. c) Probability distribution of the distance from residue 189 to the plane spanned by residues 68, 83 and 116. The curve is a sum of two gaussian distributions each shown in grey with the cumulative curve shown in green and corresponds to the distribution of the angle in the figure to the right. d) Time evolution of the shortest distance of residue 189 to the plane spanned by residues 68, 83 and 116. This distance has been found by means of Eq. (4).*

### Supporting tables with references in the main manuscript

*Table S1:* ***Summary of positively selected or migration behavior related amino acid sites of cryptochrome 4 (Cry4).*** *The Models of PamL and RateShift were executed with the whole species alignment. Coevol was run with migrant and resident passerines (PassMR).* ***ω*** *is dN/dS,* ***f*** *indicates foreground branches and* ***b*** *background branches and* ***PP*** *is the posterior probability. For the Branch-Site Model, all passerines were selected as foreground clade (Pass). For Coevol diffsel the significant amino acid is either* **+** *for an increased fitness factor or* **–** *for a decreased fitness factor in migratory passerines compared to resident passerines. The model was run six times and only sites with a consequent PP > 0.95 or < 0.05 are reported. RateShift was run with different groups: The first group consists of clades that have a passerine migratory ancestor (Ancestr. MPass, based on reconstructions with ace of R package phytools). In these clades there might be also resident and indistinct birds if they lost the migratory behavior secondarily. 525 (*) showed a significantly lower rate in only night migratory songbirds. The sites of interest show a lower evolutionary rate on the foreground branches (Ancestr. MPass) compared to background branches (RAll).*

| | **PamL (All)** | | | | | | | **Coevol (PassMR)** | | **Rateshift (All)** | | |
|---|---|---|---|---|---|---|---|---|---|---|---|---|
| | Site-Model M1a/M2a LRT: p=0.000 | | Site-Model M7/M8 LRT: p=0.000 | | Branch-Site Model Pass/NoPass; LRT: p=0.002 | | | DS3 Model diffsel | | Ancestr. MPass vs RAll | | |
| **Sites** | PP | ω | PP | ω | PP | ωf | ωb | PP | AmAcid | p-value | r.f | r.b |
| 39 | | | | | | | | | | 0.01 | 5.637 | 0.949 |
| 45 | | | | | | | | | | 0.034 | 1e-12 | 2.016 |
| 68 | 0.999 | 2.249 | 0.995 | 2.243 | | | | | | | | |
| 69 | | | | | | | | | | 0.007 | 5.574 | 1.08 |
| 83 | 1 | 2.25 | 1 | 2.25 | | | | | | | | |
| 89 | | | | | | | | | | 0.021 | 1e-12 | 2.416 |
| 90 | | | | | | | | | | 0.011 | 5.663 | 1.245 |
| 116 | 1 | 2.25 | 0.999 | 2.249 | | | | 0.957 | R+ | | | |
| 142 | | | | | | | | | | 0.033 | 2.893 | 0.302 |
| 189 | 0.9 | 2.249 | 0.998 | 2.247 | | | | | | | | |
| 193 | | | | | | | | | | 0.018 | 2.888 | 1e-12 |
| 201 | | | | | | | | | | 0.046 | 1e-12 | 1.82 |
| 202 | 1 | 2.5 | 1 | 2.25 | 0.987 | 2.26 | 1 | 0.953 | A+ | | | |
| 290 | | | | | | | | | | 0.018 | 2.805 | 1e-12 |
| 291 | | | | | | | | | | 0.018 | 1.417 | 1e-12 |
| 304 | | | | | | | | | | 0.018 | 1.52 | 1e-12 |
| 317 | | | | | | | | | | 0.049 | 1e-12 | 2.821 |
| 320 | | | | | 0.982 | 2.26 | 0.067 | | | | | |
| 330 | | | | | 0.937 | 2.26 | 1 | | | | | |
| 384 | | | | | | | | | | 0.021 | 1.537 | 1e-12 |
| 414 | | | | | | | | | | 0.005 | 3.81 | 0.489 |
| 425 | | | | | 0.992 | 2.26 | 1 | | | | | |
| 479 | | | | | | | | | | 0.017 | 1.56 | 1e-12 |
| 497 | | | | | | | | | | 0.028 | 2.93 | 0.284 |
| 510 | 1 | 2.25 | 1 | 2.25 | 1 | 2.26 | 1 | | | 0.023 | 13.68 | 4.326 |
| 512 | | | | | 0.961 | 2.26 | 1 | | | | | |
| 516 | | | | | | | | 0.034 | N- | | | |
| 517 | 1 | 2.25 | 1 | 2.25 | | | | | | | | |
| 525 | | | | | 0.974 | 2.26 | 1 | 0.981 | T+ | 0.023* | 1e-12* | 3.327* |

**Table S2: Results of codeml site and branch-site models based on the Cry4 alignment and birdtree.org tree.** The two site-model comparisons and the branch-site model testing for positive selection in passerines were significant, indicating positive selection. For M7 and M8 and branch site comparison the null distribution with 50:50 mixture of point mass 0 and $\chi_1^2$ was used (Yang and dos Reis, 2011).

| **Site-Model** | **Ln L** | **Kappa** | **W0/p** | **W1/q** | **W2/Wp** | | **Null** | **LRT** | **Df** | **p** |
|---|---|---|---|---|---|---|---|---|---|---|
| M0 | -65788.8 | 4.02 | 0.15 | | | | | | | |
| M1a | -63869.58 | 4.48 | 0.066(0.81) | 1.0(0.19) | | | | | | |
| M2a | -63825.37 | 4.53 | 0.067(0.81) | 1.0(0.18) | 1.94(0.02) | | M1a | 88.0 | 2 | 0.00<br>0 |
| M7 | -63104.36 | 3.98 | 0.23 | 1.1 | | | | | | |
| M8 | -63019.43 | 4.48 | 0.33 | 1.97 | 1.35 (0.02) | | M7 | 169.88 | 2 | 0.00<br>0 |
| **Branch-site Model** | **Ln L** | **Kappa** | **W0/p** | **W1/q** | **W2a/Wp** | **W2b/Wp** | **Null** | **LRT** | **Df** | **p** |
| Model A neutral Pass | -63835.27 | 4.48 | b: 0.06(0.79)<br>f: 0.06 | B: 1.0(0.18)<br>F: 1.0 | B: 0.06(0.03)<br>F: 1.0 | B: 1.0(0.01)<br>F: 1.0 | | | | |
| Model A selection Pass | -63829.11 | 4.5 | B: 0.07(0.8)<br>F: 0.07 | B: 1.0(0.18)<br>F: 1.0 | B: 0.07(0.02)<br>F: 2.26 | B: 1.0(0.003)<br>F: 2.26 | MA neutral | 12.34 | 2 | 0.00<br>1 |
| Model A neutral no-Pass | -63768.71 | 4.42 | B: 0.06(0.78)<br>F: 0.06 | B: 1.0(0.13)<br>F:1.0 | B: 0.06(0.07)<br>F: 1.0 | B: 1.0(0.01)<br>F: 1.0 | | | | |
| Model A selection no-Pass | -63768.71 | 4.42 | B: 0.06(0.78)<br>F: 0.06 | B: 1.0(0.13)<br>F:1.0 | B: 0.06(0.07)<br>F: 1.0 | B: 1.0(0.01)<br>F: 1.0 | MA neutral | 0 | 2 | 1 |

*Table S3: **Results of codeml site-models based on Cry1 and Cry2 alignments and birdtree.org tree.** The model comparison was significant for Cry2, indicating positive selection.*

| **Site-Model** | **Ln L** | **Kappa** | **W0/p** | **W1/p** | **W2/Wp** | **Null** | **LRT** | **Df** | **p** |
|---|---|---|---|---|---|---|---|---|---|
| M0 Cry1 | -28301.05 | 3.93 | 0.05 | | | | | | |
| M1a Cry1 | -27652.53 | 4.05 | 0.01(0.93) | 1.0(0.07) | | | | | |
| M2a Cry1 | -27652.57 | 4.06 | 0.01(0.93) | 1.0(0.07) | 1.69(0.00) | M1a | | 2 | 1 |
| M0 Cry2 | -38472.47 | 4.52 | 0.04 | | | | | | |
| M1a Cry2 | -37587.08 | 4.64 | 0.023(0.91) | 1.0(0.09) | | | | | |
| M2a Cry2 | -37579.61 | 4.64 | 0.023(0.92) | 1.0(0.08) | 2.09 (0.003) | M1a | 14.9 | 2 | 0.001 |

*Table S4: **Results of Hyphy FEL with Cry4.** Passeriformes were selected as foreground branches. The test is similar to the branch-site model of PamL. Alpha refers to synonymous and beta to non-synonymous substitutions.*

| **Site** | **Alpha** | **Beta** | **Alpha=beta** | **p** | **class** |
|---|---|---|---|---|---|
| 320 | 0.38 | 1.196 | 0.844 | 0.028 | diversifying |
| 330 | 0 | 2.224 | 1.108 | 0.000 | diversifying |
| 425 | 0.555 | 2.449 | 1.746 | 0.0006 | diversifying |
| 510 | 0.596 | 2.803 | 1.476 | 0.000 | diversifying |
| 512 | 0.604 | 1.951 | 1.347 | 0.0037 | diversifying |

*Table S5:* ***Results of Hyphy Contrast-FEL with Cry4.*** *Passeriformes were selected as foreground branches (f) and changes in selection pressure were detected between non-passerines on background branches (b) and passerines at 11 sites (with a q-value threshold of 0.02 to control for false positive discovery rate). Subs-f are substitutions mapped on foreground branches. Alpha refers to synonymous and beta to non-synonymous substitutions.*

| Site | Alpha | Beta-f | Beta-b | Subs-f | p | q-value |
|---|---|---|---|---|---|---|
| 31 | 0.635 | 0 | 1.138 | 6 | 0.000 | 0.003 |
| 45 | 0.814 | 0.13 | 1.182 | 3 | 0.000 | 0.004 |
| 135 | 2.629 | 1.599 | 0.29 | 31 | 0.000 | 0.001 |
| 181 | 0.692 | 0.184 | 1.375 | 8 | 0.000 | 0.002 |
| 255 | 0.284 | 0.302 | 1.39 | 5 | 0.000 | 0.007 |
| 320 | 0.383 | 1.218 | 0.232 | 19 | 0.000 | 0.007 |
| 490 | 0.778 | 0.295 | 1.615 | 7 | 0.000 | 0.002 |
| 510 | 0.606 | 2.813 | 0.274 | 45 | 0.000 | 0.000 |
| 517 | 0.44 | 0.438 | 2.079 | 8 | 0.000 | 0.003 |
| 523 | 0.599 | 0.456 | 0 | 14 | 0.000 | 0.016 |
| 525 | 4.392 | 2.265 | 0.549 | 43 | 0.000 | 0.017 |

*Table S6:* ***Overview of simulations carried out for the wildtype cryptochromes and the introduced mutations.***

| Cryptochrome type | Mutation introduced | Production length |
|---|---|---|
| ErCry4 | WT | 500 ns |
| ErCry4 | C317H | 200 ns |
| ErCry4 | E320K | 200 ns |
| ClCry4 | WT | 500 ns |
| ClCry4 | H317C | 200 ns |
